## Supplemental Data 1 for "HLAIIPred: Cross-Attention Mechanism for Modeling the Interaction of HLA Class II Molecules with Peptides"

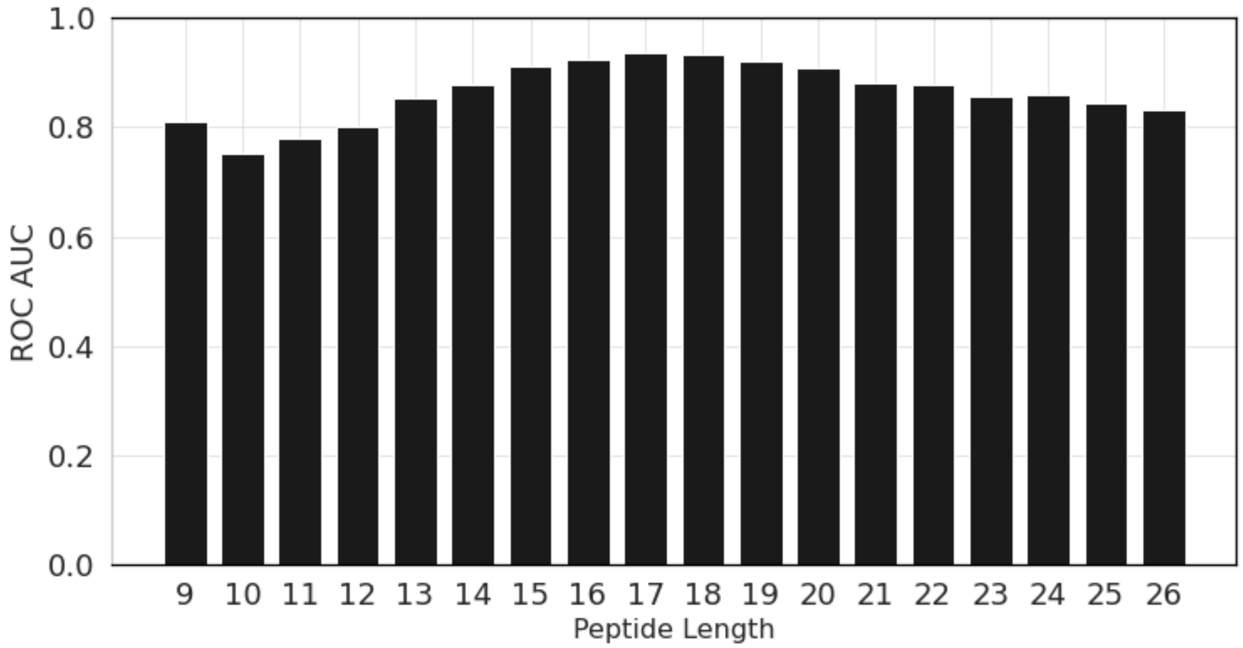


*Supplementary Figure 1 depicts the performance of HLAIIPred in terms of ROC-AUC for different peptide lengths in the test set.*

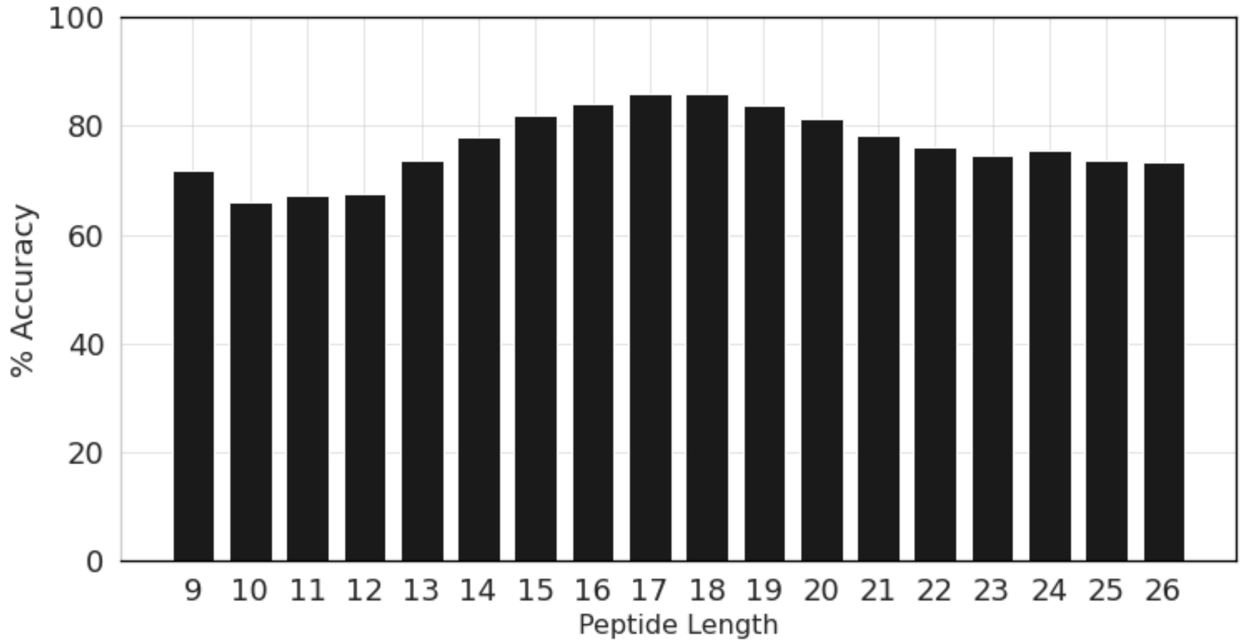


*Supplementary Figure 2 depicts the performance of HLAIIPred in terms of Accuracy(%) for different peptide lengths in the test set.*
